# Chaos in small microbial communities

**DOI:** 10.1101/2021.09.06.459097

**Authors:** Behzad D. Karkaria, Angelika Manhart, Alex J. H. Fedorec, Chris P. Barnes

## Abstract

Predictability is a fundamental requirement in biological engineering. As we move to building coordinated multicellular systems, the potential for such systems to display chaotic behaviour becomes a concern. Therefore understanding which systems show chaos is an important design consideration. We developed a methodology to explore the potential for chaotic dynamics in small microbial communities governed by resource competition, intercellular communication and competitive bacteriocin interactions. We show that we can expect to find chaotic states in relatively small synthetic microbial systems, understand the governing dynamics and provide insights into how to control such systems. This work is the first to query the existence of chaotic behaviour in synthetic microbial communities and has important ramifications for the fields of biotechnology, bioprocessing and synthetic biology.

## 1 Introduction

Chaos can be defined as deterministic behaviour that displays aperiodic orbits and sensitivity to initial conditions [1]. Infinitesimally small differences in initial conditions of a chaotic system will become amplified over time, making forecasting and prediction of behaviour impossible [2]. Despite being deterministic, chaotic systems possess an inherent uncertainty due fact that we can never describe the initial conditions of a system in sufficient detail. Building systems which behave in a predictable and repeatable manner is essential across fields invested in engineering biology and its applications. Evidence from studies of neural networks suggests the increasing probability of chaotic behaviour as the number of dimensions in the network grow [3, 4, 5]. Therefore we might expect opportunities for unpredictable behaviour to become more probable as we try and implement larger synthetic communities, or edit existing networks such as the human gut microbiome. Steps to date have not been taken to investigate the existence of chaos in small synthetic microbial networks. A long-term goal of engineering biology is to create truly scalable and robust synthetic microbial communities [6, 7]. Therefore understanding and evaluating the possibility of chaotic behaviour in a system becomes an important consideration.

Observations of chaotic behaviour in biological systems have been reported. A three species system containing one predator and two prey species has been demonstrated to produce chaotic behaviour, with dilution rate a key parameter in enabling aperiodic behaviours [8]. An eight year study of a planktonic food web measured chaotic behaviours, resulting in subpopulation abundance predictability being limited to 15-30 days, despite constant external conditions [9]. These experimental examples demonstrate that a low number of species are capable of producing chaotic behaviour and are therefore unpredictable.

In order to predict chaotic behaviour in synthetic microbial communities, we need to develop models that capture interactions between different community species. Generalised competitive Lotka-Volterra equations (gLV) have previously been used to model pair-wise interactions and infer inter-species relationships [10]. However in other circumstances, gLV models provide an incomplete description of interactions we expect to find in microbial communities. They are unable to capture the existence of chaos in three species networks [11]. Furthermore, gLV models have failed to predict community formation from pairwise interactions in microbial communities [12]. gLV models lack dynamics that occur with the accumulation and depletion of extracellular species, which can be important for predicting the true dynamics of a community [13]. Modified Lotka-Volterra equations produce chaotic behaviour in predator-prey systems by including time-delayed feedback [14, 13], or in one predator two prey systems, by adding dampening effects [15]. While these abstractions are suitable in some circumstances, using them to inform gene regulation networks and community design can be difficult. By modelling the intermediates involved in competitive interactions we can include experimentally measurable mechanisms and parameters. In previous work, we have modelled quorum sensing (QS) to regulate bacteriocin expression and engineer inter-population interactions. These methods allowed us to tune experimental parameters of an existing two strain system [16], and predict the most promising topologies for producing stability in two and three strain systems [17].

The existence of chaos in dynamical models and identification of chaotic parameter space can be identified using various optimisation techniques. The unscented Kalman filter has previously been used to investigate chaos in electrical circuits and biological systems, obtaining parameters yielding chaos [18]. Simulated annealing has been applied to finding chaotic parameters in four species standard Lotka Volterra models [19]. Evidence also suggests that perturbation of system parameters can be used to drive systems towards or away from chaotic attractors [20]. The possibility of chaos in synthetic microbial communities, to our knowledge, has not been previously considered.

## 2 Results

### 2.1 Searching for chaos in microbial community models

In previous work we developed a model framework to describe QS regulated bacteriocin interactions in a three strain model space, and predicted topologies that form stable communities [17]. Here we use this same model space to investigate the existence of chaos in three strain synthetic microbial communities.

Figure 1a shows the pipeline we developed to search for chaos in synthetic three strain systems. The initial model space describes an enumeration of possible combinations of bacteriocin and QS systems. Prior parameter distributions describe the range of characteristics for the different parts (Table 1). We expected the existence of chaos to be sparse in this three strain model space, and therefore computationally expensive to explore. Approximate Bayesian Computation Sequential Monte Carlo (ABC SMC) is a method that can be used for model selection and parameter inference in dynamical systems [21]. The definitions we use to classify oscillatory and chaotic behaviours are described in *Methods*. We also define an extinction threshold of 10^−5^, if a strain population falls below this it is classified as extinct. In order to narrow down the search, we first performed ABC SMC for an oscillations objective. Oscillations are a known route to chaos [1]. Although this is not an exhaustive search, we assumed it to be sufficient for prioritising in our search for chaos. We identified 117 models capable of producing oscillations, these models were then considered for the chaos objective (Figure 1a).

**Table 1:** Prior distributions for both two and three strain systems are sampled uniformly between the min and max values listed below. Constant parameters have the same min and max value

| Parameter / State variable | Description | Prior (min) | Prior (max) | Units | Citation |
| --- | --- | --- | --- | --- | --- |
| Parameters |  |  |  |  |  |
| $C_N$ | OD to cell number scaling factor | 1e9 | 1e9 | None | N/A |
| $C_B$ | Microcin scaling factor | 1e-9 | 1e-9 | None | N/A |
| $C_A$ | QS scaling factor | 1e-9 | 1e-9 | None | N/A |
| $D$ | Dilution rate | 0.01 | 0.5 | $h^{-1}$ | N/A |
| $K_{A_y B_z}$ | Half maximal QS promoter activation/repression from $A_y$ to $B_z$ | 1e-9 | 1e-6 | $M$ | [40] |
| $K$ | Monod's half saturation constant | 3.9e-5 | 3.9e-5 | $M$ | [41] |
| $K_\omega$ | Half saturation killing constant | 1e-7 | 1e-6 | $M$ | [42, 43] |
| $S_0$ | Substrate concentration of input media (0.4% glucose) | 0.02 | 0.02 | $M$ | M9 media |
| $\gamma$ | *E. coli* substrate yield | 1e11 | 1e11 | cell $M^{-1}$ | [44] |
| $k_{A_y}$ | Production rate of AHL per cell | 1e-22 | 1e-15 | $M h^{-1}$ | [45] |
| $KB_{max} Z$ | Maximal expression rate of microcin | 1e-22 | 1e-15 | $M h^{-1}$ | [46] |
| $\mu_{x_{max}}$ | Maximum growth rate | 0.4 | 3 | $h^{-1}$ | [47, 48] |
| $n_z$ | Hill coefficient AHL induced expression | 1 | 2 | $M$ | [40] |
| $n_\omega$ | Hill coefficient for killing | 1 | 2 | $M$ | [40] |
| $\omega_{max}$ | Maximum rate of bacteriocin killing | 0.5 | 2.0 | $M^{-1} h^{-1}$ | [49, 42, 43] |
| Initial state variable |  |  |  |  |  |
| $N$ | OD of strain | 0.01 | 0.5 | OD | N/A |
| $S$ | 0.4% glucose concentration | 0.02 | 0.02 | $M$ | N/A |
| $B$ | Microcin concentration | 1e-81 | 1e-81 | $M C_B$ | N/A |
| $A$ | QS concentration | 1e-10 | 1e-10 | $M C_A$ | N/A |

**Figure 1:**
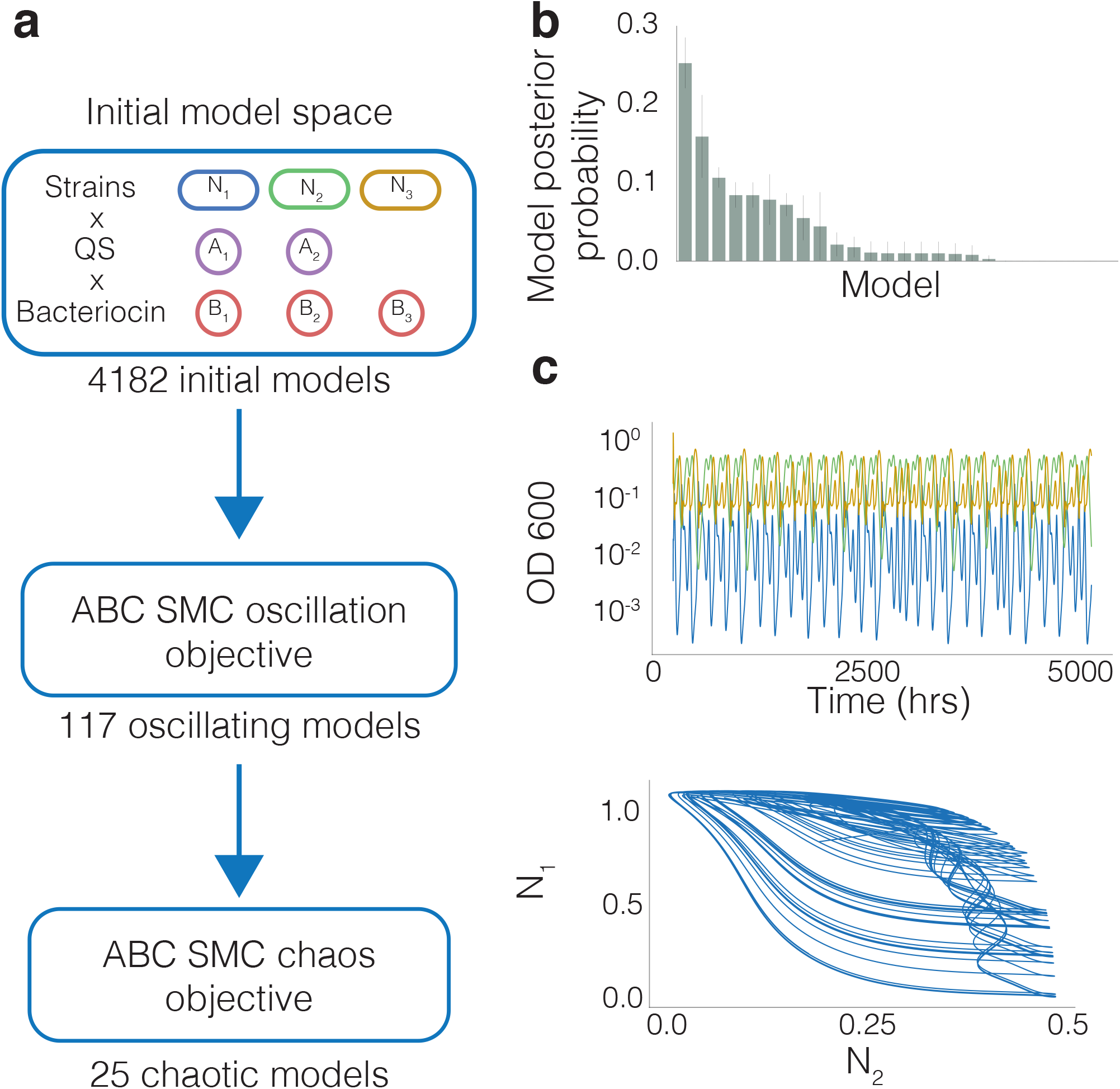
Overview of the pipeline for identifying chaotic topologies **a** By combining engineering options in different combinations, we generate 4182 models that form our initial model space. We then perform ABC SMC for an oscillatory objective which yielded 117 models that were capable of producing oscillations. These form the prior model space for the chaos objective. **b** The barchart shows the probability of models for the chaotic objective. The error bars represent the standard deviation. **c** An example time series representative of the dataset; it shows sustained, nonrepetitive oscillatory behaviour for the three species community.

The chaos objective is defined by calculating the maximal Lyapunov exponent (λ_1_). We calculate λ_1_ by initialising two nearby orbits and measuring their divergence or convergence over the course of a simulation (Methods). λ_1_ < 0 corresponds to linear stability, λ_0_ = 0 corresponds to periodic oscillations, and λ_1_ > 0 corresponds to chaos. Due to the limited time frame from which calculate the Lyapunov exponent, oscillations are unlikely to give rise to precisely λ_1_ = 0. By running ABC SMC for an objective of λ > 0 and manually inspecting the trajectories in this population we determined a threshold of λ_1_ > 0.003, above which we are confident only chaotic behaviour exists, and below which only periodic oscillatory behaviour exists. Performing ABC SMC for the chaotic objective (λ > 0.003), we identified 25 models that produced chaotic behaviour. The posterior probabilities of the models are shown in Figure 1b. Figure 1c shows a representative chaotic trajectory, demonstrating aperiodic non-repeating behaviour, satisfying the qualitative features of chaos.

### 2.2 Properties of chaotic models

We next explored some of the properties of chaotic topologies we identified using ABC SMC. Figure 2a shows the top performing models when subsetting for complexity, based on the number of parts expressed. *m*_850_ contains four expressed parts and possess the highest posterior probability for chaotic behaviour. Systems containing fewer parts all had a posterior probability of zero. As complexity increases to five and six parts (*m*_3177_ and *m*_2547_), the posterior probability decreases. The classic debate on the complexity-stability relationship in theoretical ecology is likely highly dependent on the nature of the biological interactions involved [22, 23], but here we see some evidence for a peak in the probability of chaotic behaviour at four parts. This is in contrast to our previous findings, where system stability increased with the number of parts [17].

**Figure 2:**
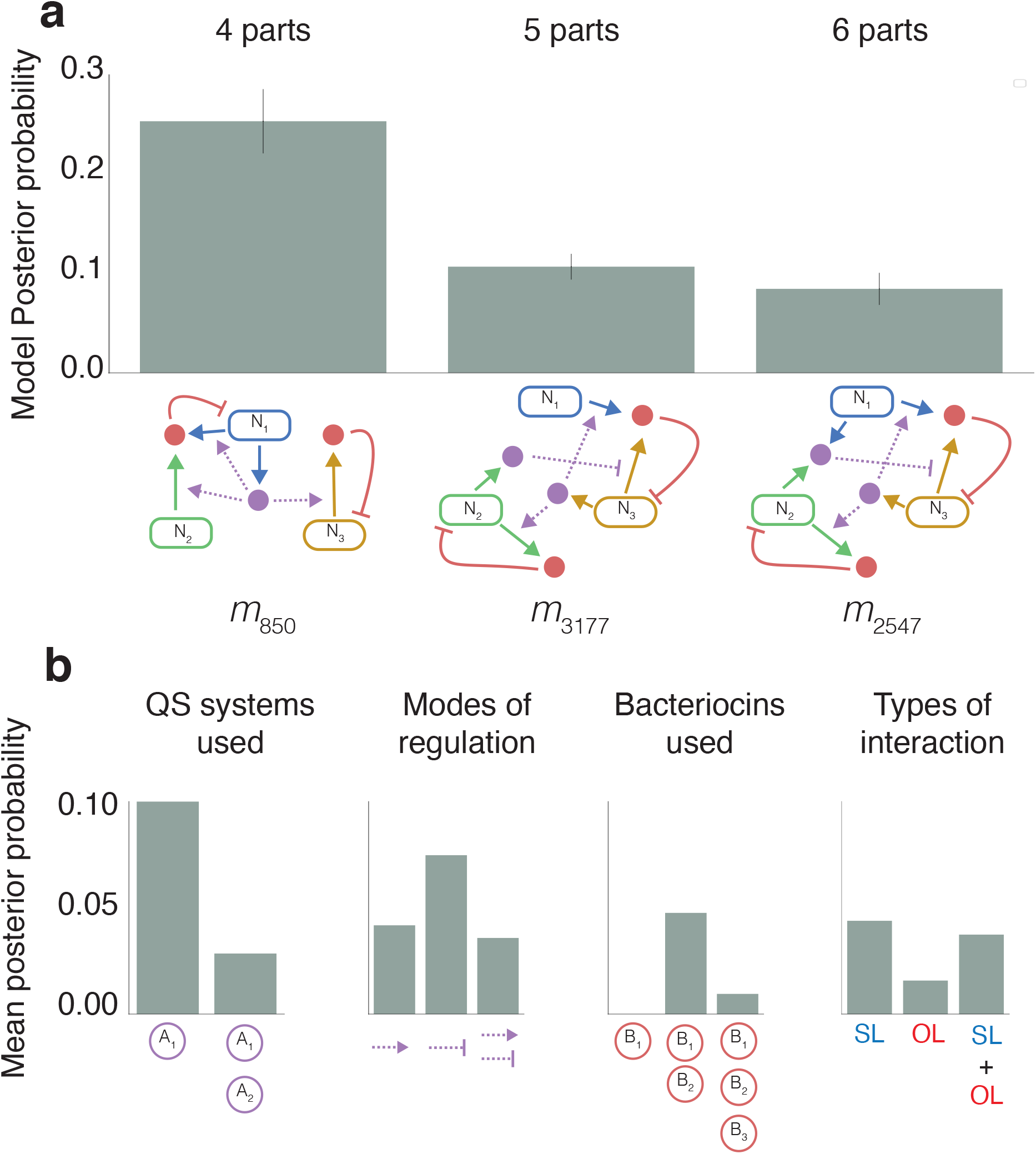
Topologies and properties associated with chaotic behaviour **a** Shows the models with highest posterior probability when subsetted for number of parts expressed, in order of increasing complexity (4, 5 and 6 expressed parts). The bar chart shows the mean model posterior probability across three experiments, represented by the scatter points, the error bars indicate the standard deviation. **b** Comparison between average posterior probabilities with different properties. In order from left to right, the barcharts compare: The number of QS systems used, the modes by which QS regulates bacteriocin expression (positive, negative or both), the number of bacteriocins used, and systems containing self-limiting (SL), other-limiting (OL) or SL and OL interactions.

Figure 2b provides summaries of how different parts contribute to chaotic behaviour in the three strain models. We can see that one QS system and positive regulation of bacteriocin is strongly favoured for producing chaos. This ensures all system bacteriocins are regulated in tandem. Expression rates are all dependent upon the same QS, resulting in stronger negative or positive correlations defined by the mode of regulation. Two bacteriocin systems also dominate the model posterior. Bacteriocin interactions can be categorised as either self-limiting (SL), whereby the strain is inhibited by the bacteriocin it produces, or other-limiting (OL) where a strain is inhibited by a bacteriocin produced by a different strain. Both SL only and a combination of SL and OL interactions are associated with producing chaotic behaviour. These observations are interesting in comparison to other work on ecological systems. Cooperative interactions were previously found to give rise to unstable systems, whereas competition was more indicative of stability [24]. The same effect might occur here in systems with one QS, rather than two, as the system would be expected to have increased correlation. While chaotic behaviour appears to be very different from linear stability, both behaviours share the necessity for coexistence. This may explain why we see tendencies for topologies to share a mixture of stability associated SL interactions, and instability associated OL interactions. We also find models with three bacteriocins, and hence higher suppression of growth, have a low posterior probability for chaos.

### 2.3 Parameter importance for chaos

The model with the highest posterior probability for chaotic behaviour was *m*_850_, the topology is shown in Figure 3a. It consists of a single QS system, produced by *N*_1_, that positively regulates two bacteriocins. *B*_1_ is produced by *N*_1_ and *N*_2_ but it inhibits the growth of *N*_1_ only. *B*_2_ is produced by *N*_3_ and inhibits the growth of *N*_3_ only. The system in total consists of four expressed parts. *m*_850_ also ranked highly for the oscillatory objective, ranking 3rd out of the initial 4182 models. This presents an interesting problem whereby a model that has promising use as an oscillator also has a high potential to produce chaos, relative to other candidate models. Identifying the parameters and initial conditions important for differentiating between chaotic and oscillatory behaviour gives us insight into how to control this behaviour when constructing genetic circuits or selecting chemostat settings.

**Figure 3:**
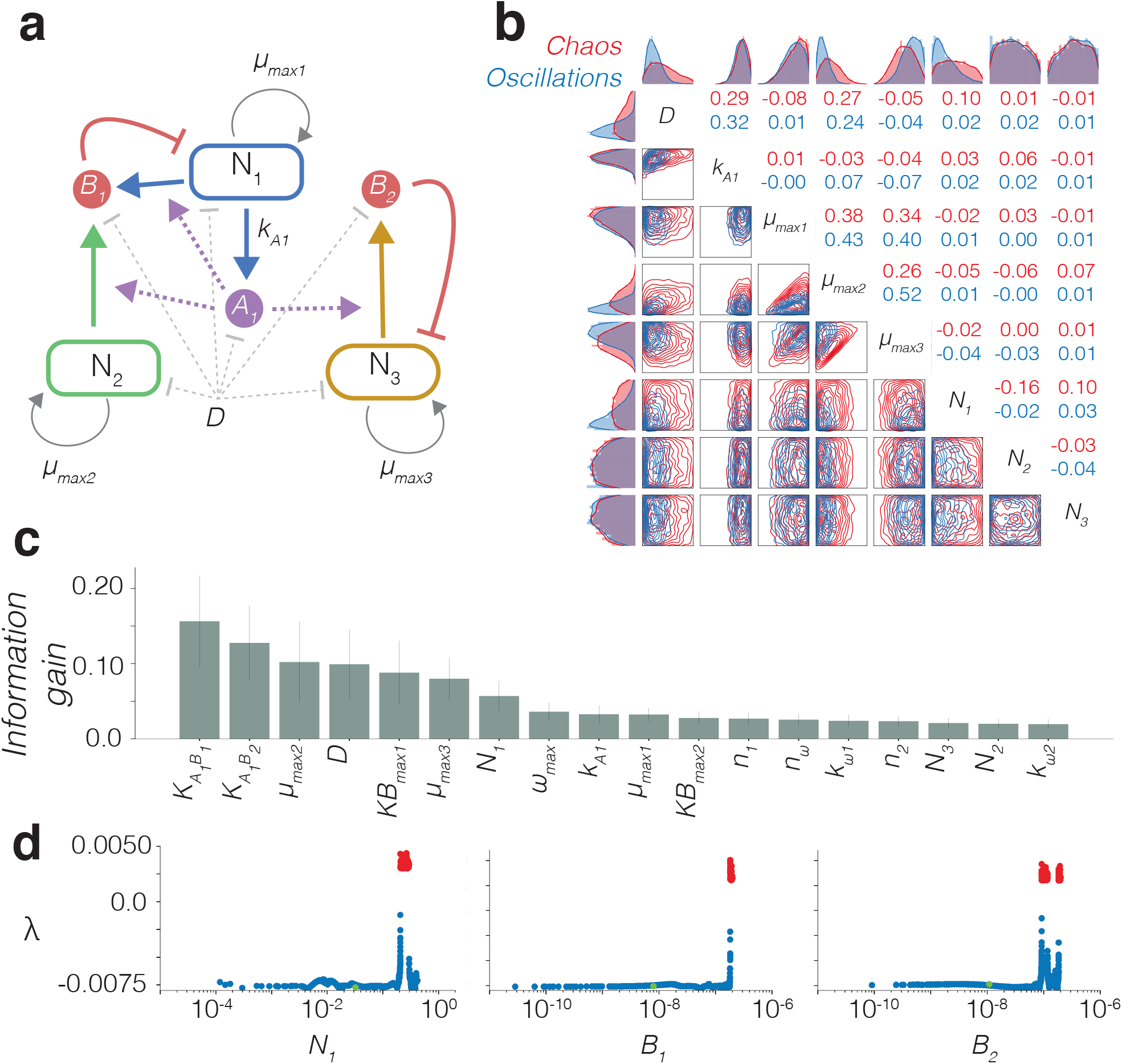
Examining chaos in *m*_850_. **a** Topology of *m*_850_ with key parameters labelled. *k*_*A*1_ is the rate of QS molecule production, *KB*_*max*1_ and *KB*_*max*2_ are the maximal expression rates of bacteriocins *B*_1_ and *B*_2_ respectively. **b** Posterior parameter distributions of *m*_850_ for chaos (red) and oscillatory (blue) objectives for key parameters in system design. The borders show 1D posterior distributions for each parameter and the off-diagonal element the 2D posterior marginals. **c** Feature importance calculated using random forest regression. The information gain (bits) is calculated as an average of the reduction in entropy across all trees in the forest (2000 trees). The error bars indicate the standard deviation of the entropy for each feature across all trees. **d** Sensitivity analysis of a chaotic input vector with chaotic region in red. Green dots refer to the identified stable steady state. The fixed parameter values are shown in Table 2

As a first step, we analyzed the model to quantify the possible steady states and basins of attraction. Our analysis gave analytical conditions for the existence and stability for complete extinction and for single strain survival (See Methods). For three-strain co-existence, we find the following necessary conditions:

**Table 2:**
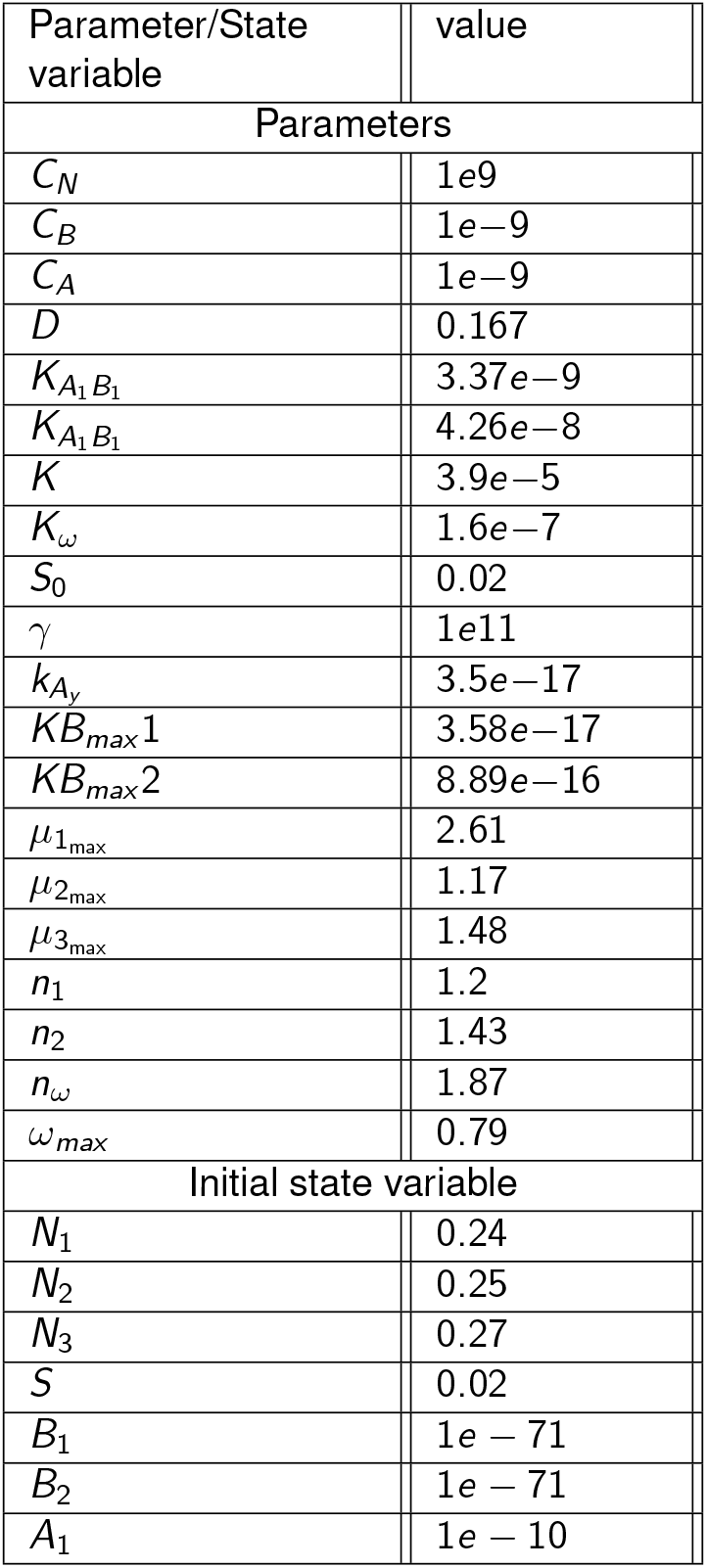
Fixed parameters used in Figures 3d and Figures 4.

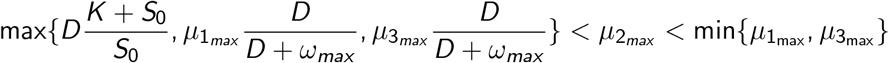

This shows that for three-strain co-existence, the maximal growth rate of *N*_2_ has to lie between certain upper and lower bounds. In particular, it has to be smaller than the maximal growth rate of *N*_1_ or *N*_3_. We can see from the topology of *m*_850_ (Figure 3a) that the growth of *N*_2_ is not limited by any bacteriocin, therefore the only limitation on growth comes through resource competition. If *N*_2_ had a higher growth rate than *N*_1_ or *N*_3_, it would out compete these strains and cause an extinction event.

We then wanted to explore the most important parameters that separate oscillatory and chaotic behaviours in *m*_850_ only. We refer to a set of parameters and initial conditions as an input vector. Using ABC SMC, we performed parameter inference on *m*_850_ for the chaotic and oscillatory objectives, generating 3750 input vectors for each objective. We can use this dataset of labelled input vectors to understand the importance of individual parameters, initial conditions and nearby steady states.

Figure 3b shows multivariate parameter distributions for the oscillator and chaotic objectives for the experimentally accessible parameters. The dilution rate (*D*) is a directly controllable parameter of the chemostat. The production rate of *A*_1_ (*kA*_1_) can be tuned by using an inducible promoter to control expression of the AHL synthase species. Strain maximal growth rates (*µ*_*max*1_, *µ*_*max*2_, *µ*_*max*3_) can be controlled by using different base strains or through the combined use of auxotrophic strains and defined media. Finally, the initial population densities (*N*_1_, *N*_2_, *N*_3_) can easily be set when inoculating the initial culture. Divergence between two parameter distributions indicates its importance in differentiating between the two objectives. We can see that the oscillatory objective distributions for *D, N*_1_ and *µ*_*max*2_ are all constrained towards lower values relative to the prior. However, for all these distributions we can see that the chaotic and oscillatory regions overlap. This again implies that chaotic and oscillatory behaviour exist close to one another in parameter space, and highlights the multidimensional nature that determines the behaviour.

To further investigate the importance of parameters and initial conditions we trained a random forest classifier model using the input vectors as features. This classifier model was able to classify the test set with a ∼90% accuracy (Methods, Figure 6). Figure 3c shows the average information gain across all decision tree classifiers in the forest for all free parameters. This can be used as an indicator of feature importance in correctly classifying an input vector. 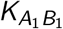 and 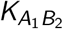 describe the concentration of *A*_1_ required to produce half-maximal repression of bacteriocins *B*_1_ and *B*_2_ respectively. While the feature importance indicates these parameters are the most important, they are more difficult to tune compared with other parameters in this system. The error bars indicate the variability in the importance of a feature across all trees in the forest. Large error bars suggest single features are not essential for classification, and that redundancy exists between the features used [25].

From the set of chaotic input vectors, we used numerical methods to identify nearby steady states by changing the initial state of the system. Figure 3d shows the sensitivity analysis of a chaotic input vector, with a nearby three-strain stable steady state shown in green. Starting from the stable steady state (green), we perturbed the initial species values of either *N*_1_, *B*_1_ or *B*_2_ individually. The plots show how changing these initial states yields different Lyapounv exponents, highlighting the chaotic region in red. The range of Lyapunov exponents shown in Figure 3d suggest that by changing the initial conditions only we are able to produce a range of different behaviours. Perturbing *N*_2_, *N*_3_ or *A*_1_ did not produce chaotic behaviour. It is interesting that the initial state of *N*_1_ as the *A*_1_ producing strain appears to more important whereas the initial concentration of *A*_1_ itself is not.

### 2.4 Exploring the parameters in the transition to chaos

Being able to move a system from a chaotic state to a fixed point could be important in a bioprocess control scenario so we explored this in more detail. Previous studies have frequently identified the bioreactor dilution rate as an important parameter for transitioning between different population dynamics [26, 27, 28]. Figures 3b strongly indicated *D* to be important for defining chaotic behaviour. We previously identified the QS production rate, *k*_*A*1_ and the dilution rate, *D*, as important parameters for transitioning between co-existence and extinction states [16]. We hypothesised that the antagonistic effect of *k*_*A*1_ to *D* would make it also make it a useful parameter for controlling population behaviour.

First, we took an input vector known to produce chaotic behaviour and randomly sampled new values for *k*_*A*1_ and *D* from the prior and calculated the Lyapunov exponent of the new input vector. Figure 4a shows the results where filled colour indicates the maximal Lyapunov exponent calculated at each grid reference. The grid outline indicates the classification range and the red grid region of Figure 4a shows the chaotic region. As can clearly be seen, changing *D* and *k*_*A*1_ affects the Lyapunov exponent. The bifurcation diagrams in Figure 4b and c for *k*_*A*1_ and *D* respectively, illustrate the antagonistic transitions in behaviour that occur when changing the two parameters. Figure 4a and b show transitions through one strain extinctions (*N*_*x*_ < 10^−5^, stable steady state, oscillations and chaotic behaviour. Figure 4a and c both show that increasing *k*_*A*1_ results in transitions from stable co-existence, through oscillations and then to chaos, followed abruptly by an extinction event. Figure 4b and c both show that low a lower dilution rate is associated with chaos; increasing the dilution rate reduces instability to produce oscillations, which abruptly transitions to a stable extinction state.

**Figure 4:**
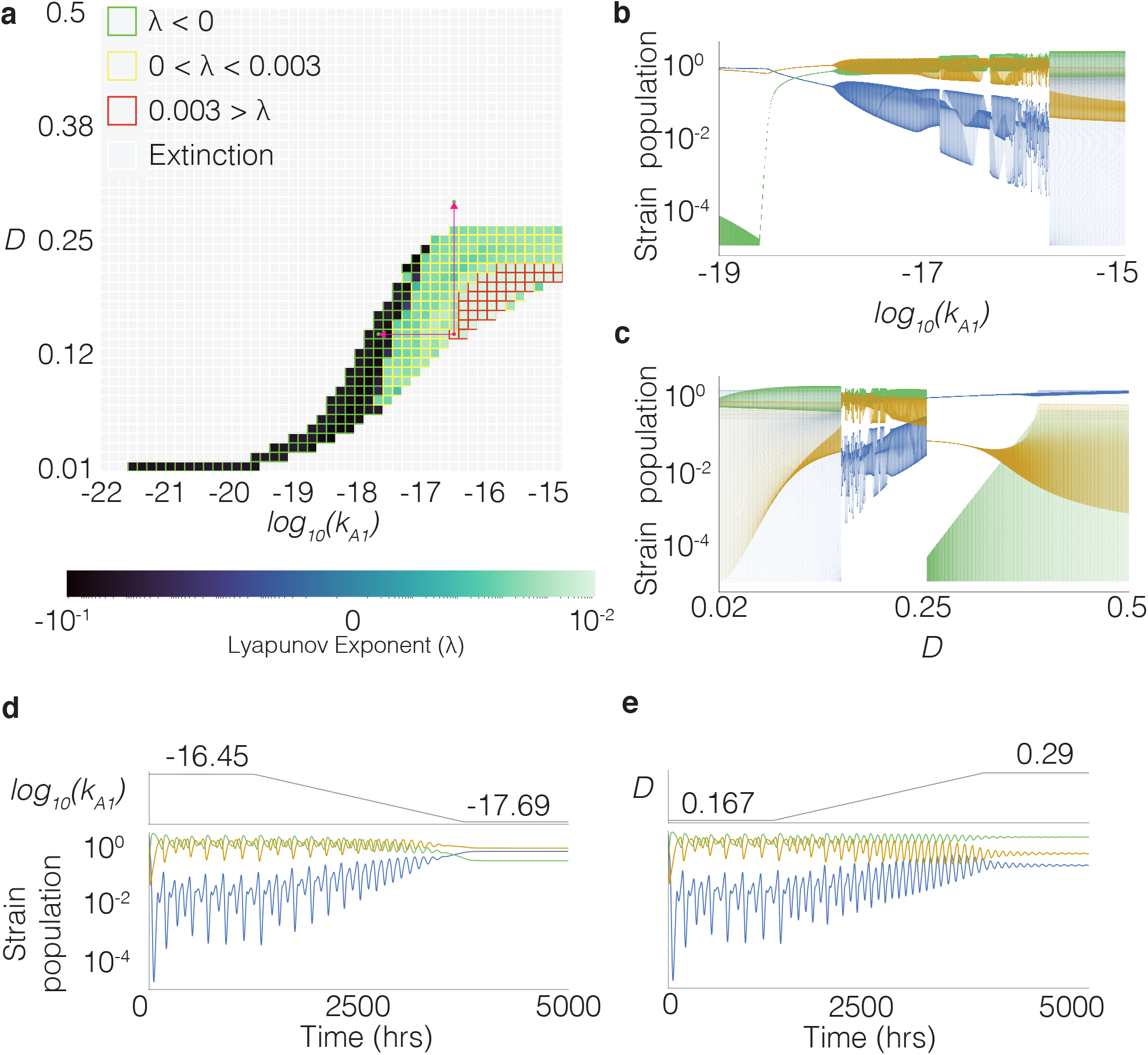
Parameters *k*_*A*1_ and *D* can be tuned to control transitions between chaotic, oscillatory and stable states. The fixed parameter values are shown in Table 2. **a** Map showing how different combinations of *k*_*A*1_ and *D* change population behaviour. The grid fill colour corresponds to the maximum Lyapunov exponent measured, the grid outlines indicated the approximate classification where green is stable, yellow is oscillatory, red is chaotic and white is extinction. **b** Bifurcation diagram showing the community states visited for different values of *k*_*A*1_. **c** Bifurcation diagram showing the community states visited for different values of *D*. **d** Real-time ramp down tuning of *k*_*A*1_, moving the system from a chaotic state to a stable steady state. **e** Real-time ramp up tuning of *D*, moving the system from a chaotic state to a stable steady state.

In a bioreactor control scenario it is interesting to understand if a community could be switched between states in real time. Figures 4d and e show how this is possible by modifying *k*_*A*1_ and *D* respectively. The red arrows on Figure 4a indicate the position of the single start point and two end points in these real-time transitions. It’s important to note that when ramping up the dilution rate in real-time, we reach stable steady state in a region that would not be obtainable with a fixed dilution rate.

## 3 Discussion/Conclusions

We developed a novel methodology to explore parameter regions that give rise to chaotic dynamics. We applied it to the exploration of chaotic dynamics in synthetic microbial communities and found a high prevalence of such dynamics in these systems. This work is the first to query the existence of chaotic behaviour in synthetic microbial communities. We show that we can expect to find chaotic states in relatively small synthetic microbial systems, which has important ramifications for the field.

We expect it will become increasingly important to consider the location of chaotic attractors in parameter space as the microbial communities we build or interact with become more complex. These methods can easily be applied to parametrise different models. It would be interesting to compare the existence of chaotic attractors in systems that use toxin-antitoxin systems [29], combination of cooperative and competitive interactions [30], or mutualistic only interactions [31]. Full scale metabolic models contain a large number of linear reactions [32], they can be combined to describe microbial communities and used to model industrial bioprocesses [33, 34]. Given the high dimensional nature of metabolic networks, it would be interesting to investigate whether these models yield chaotic behaviour in small community networks.

To conclude, we have developed methods for identifying chaotic parameter regions using ABC SMC. Although chaotic attractors are generally thought to be sparse in low dimensional systems, we have shown their existence in realistic synthetic microbial systems. They may also exist in close proximity to stable steady state regions. This work demonstrates that deterministic chaos will be an important factor in microbial community design and should be studied in much more detail.

## 4 Methods

### 4.1 Model space

Models are generated from a set of parts, which are expressed by different strains in the system. We represent an expression configuration through a set of options. We define the options for expression of *A* in each strain, where the options are not expressed, expression of *A*_1_, and expression of *A*_2_ (0, 1 and 2). We define the options for expression of bacteriocin, which for the two strain model space includes no expression, expression of *B*_1_ or expression of *B*_2_ (0, 1, and 2). For the three strain model space, this includes includes no expression, expression of *B*_1_, expression of *B*_2_ or expression of *B*_3_ (0, 1, 2 and 3 respectively). Lastly we define the mode of regulation for the bacteriocin, which can be either induced or repressed (0 and 1). This is redundant if a bacteriocin is not expressed.

Two strain model space:

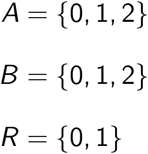

Three strain model space:

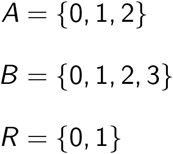

This enables us to build possible part combinations that can be expressed by a population. Let *P*_*c*_ be a family of sets, where each set is a unique combination of parts.

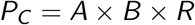

Each strain in a system can be sensitive to up to one bacteriocin. Let *I* represent the options for strain sensitivity. In the two strain model space, the options are insensitive, sensitive to *B*_1_ or sensitive to *B*_2_ (0, 1 and 2 respectively). In the three strain model space, where the options are insensitive, sensitive to *B*_1_, sensitive to *B*_2_ or sensitive to *B*_3_ (0, 1, 2 and 3 respectively).

Two strain model space:

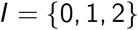

Three strain model space:

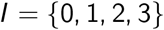

Each strain is defined by it’s sensitivities, and expression of parts. Let *P*_*E*_ be all unique engineered strains:

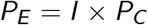

Which can be combined to form a model, yielding unique combinations in two strains and three strains:

Two strain model space:

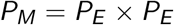

Three strain model space:

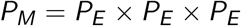

Finally, we use a series of rules to remove redundant models. A system is removed if:

1. Two or more strains are identical, concerning bacteriocin sensitivity and combination of expressed parts.
2. The QS regulating a bacteriocin is not present in the system.
3. A strain is sensitive to a bacteriocin that does not exist in the system.
4. A bacteriocin exists that no strain is sensitive to.

This cleanup yields the options which are used to generate ODE equations for a system.

### 4.2 System equations

State variables in each system are rescaled to improve speed of obtaining numerical approximations. *N*_*X*_ describes the concentration of a strain, *B*_*z*_ describes the concentration of a bacteriocin and *A*_*y*_ describes the concentration of a quorum molecule. *C*_*N*_, *C*_*B*_ and *C*_*A*_ are scaling factors:

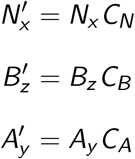

Each model is represented as sets where ℕ defines the number of strains, 𝔹 defines the set of bacteriocins and 𝔸 defines the set of QS systems. The following differential equations are used to represent each model.

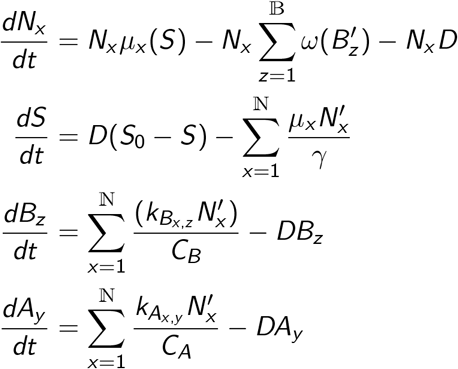

Growth modelled by Monod’s equation for growth limiting nutrient, 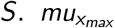 defines the maximal growth rate of the strain and *K*_*X*_ defines the concentration of substrate required for half-maximal growth.

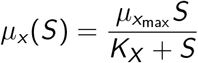

Killing by bacteriocin is defined by 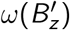, where *ω*_*max*_ defines the maximal killing rate which is set to 0 if the strain is insensitive. *K*_*ω*_ defines the concentration at which half-maximal killing occurs.

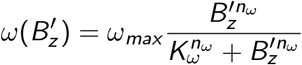

Induction or repression of bacteriocin expression by QS, is defined by *k*_*B*_ (*z, y*), where *z* defines the bacteriocin being expressed and *y* defines the quorum molecule regulating its expression. *KB*_*max*_ *z* is the maximal expression rate of the bacteriocin and 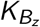 is the concentration of quorum molecule at which bacteriocin is half-maximal. *n*_*z*_ defines the cooperativity of the AHL binding.

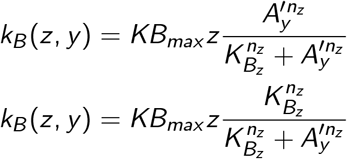

### 4.3 Software packages and simulation settings

The ABC SMC model selection algorithm was written in python using Numpy [35], Pandas and Scipy [36]. ODE simulations were conducted in C++ with a Rosenbrock 4 stepper from the Boost library [37]. All simulations use an absolute error tolerance of 1*e*−9, and relative error tolerance of 1*e* − 4. Non negative matrix factorisation was conducted using Scikit-learn [38]. Simulations were conducted for 5000hrs, and were stopped early if the population of any strain fell below 1*e* − 5 (extinction event). Simulations with an extinction event have distances set to maximum in order to prevent excessive time spent simulating collapsed populations.

### 4.4 Oscillatory population dynamic objective

We define the oscillatory population dynamic using three summary statistics for each strain. First, we use Fourier transform of the population signal to find the maximum frequency, *f*, and convert this to the period, T.

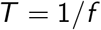

We set a minimum period of *t/*2 where *t* is the simulation time, giving us 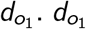. Any simulations in which 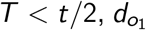 is set to 0, this distance ensures that all we have frequencies of oscillations that are on a scale relevant to the time period being measured. It was found that using the signal frequency alone resulted in acceptance of many simulations with very small oscillations, or simulations that rapidly dampen. We therefore generated two additional distances that account for oscillation amplitudes to select for sustained oscillations only. We can define the number of expected peaks in the simulation, *p*.

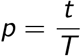

Peaks in the trajectory are identified by changes from a positive gradient to a negative gradient, and troughs via changes from negative gradient to positive gradient. The peak-to-peak amplitudes are calculated by differences between consecutive peaks and troughs. *A*_*K*_ is the number of amplitudes above the threshold, 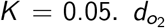 is the difference between the number of expected oscillations in the simulation, and the count of above threshold oscillations. Because incomplete oscillations at the time the simulation ends can impact the distance measurement, we set a lenient final distance threshold for 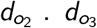 compares the final amplitude *A*_*F*_ in the simulation to the threshold. We set 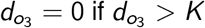. If any strain falls below an OD of 10^−5^, the population is deemed extinct and the particle rejected.

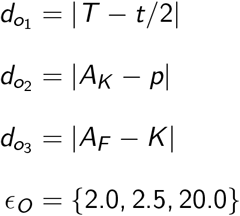

### 4.5 Maximal Lyapunov exponent calculation

Lyapunov exponents can be used to measure chaotic behaviour; they describe the average exponential rate of divergence between two near trajectories of a dynamical system. The maximal Lyapunov exponent, λ_1_, can be used as determinant of chaotic behaviour. Using a method described by Sprott et al. [39], We evolve two nearby orbits and measure their average rate of separation. This directly investigates whether small changes to an initial state will produce a disproportionate separation. By periodically readjusting the distance of divergence after each time step we measure separation across a period of time, preventing a single event dominating subsequent states (Figure 5). The method is described in full by Algorithm 1. For all simulations we generate nearby orbits by perturbing one of the strain initial strain densities by *d*_0_ = 10^−10^. All simulations use a transient time equivalent to the first 10% of the time series.

**Figure 5:**
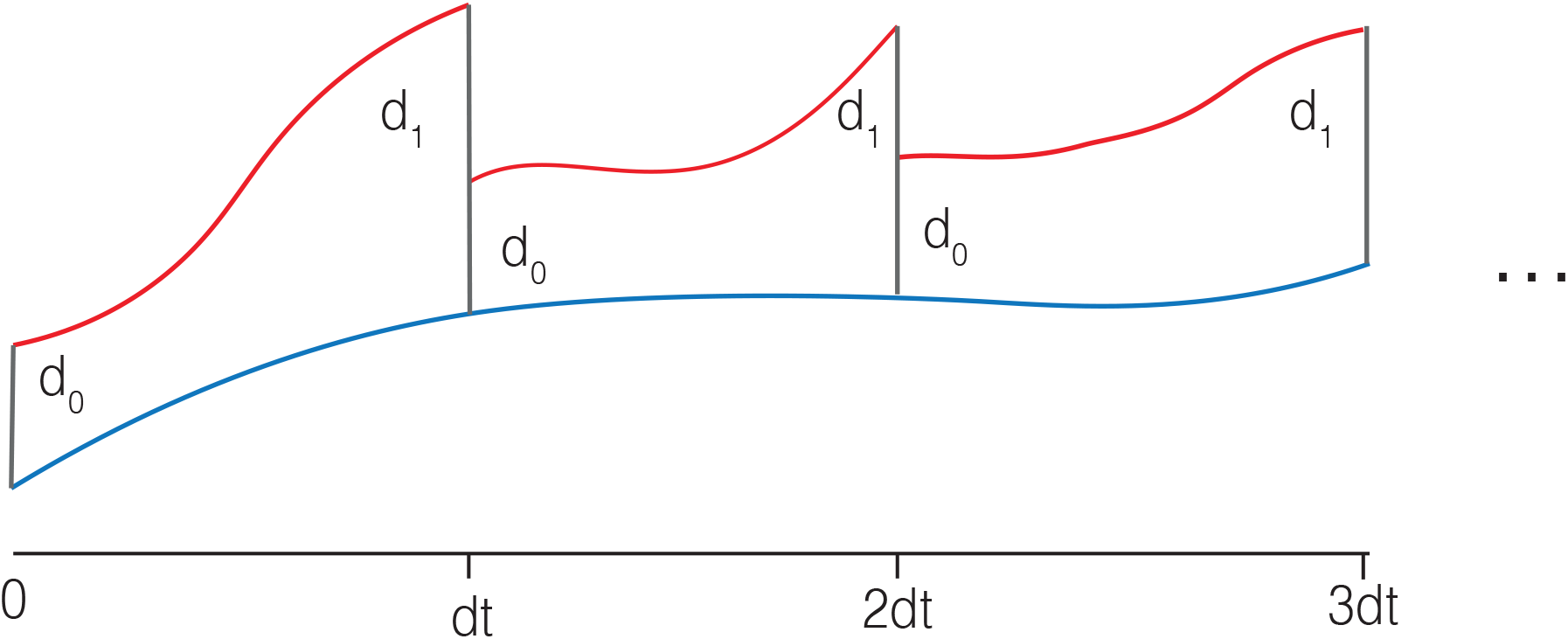
Illustration of dual-orbit algorithm used to calculate the λ_1_. Two orbits with an initial state separation of *d*_0_ are followed. After each time step measure the separation, *d*_1_, is measured. The perturbed orbit (red) is readjusted to prevent excess separation. The average rate of separation between the two orbits corresponds with the λ_1_.

### 4.6 Chaos population dynamic objective

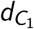 is the only distance for the chaotic objective. If 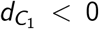, the particle is rejected. The final distance threshold, *ϵ*_*C*_, is equivalent to all λ_1_ > 0.003.

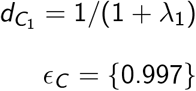

For each sampled particle a prescreening process was performed to minimise time spent conducting the more computationally time consuming dual-orbit method. Simulations in which an strain fell below 1e-5 were rejected. The number of oscillations with an amplitude greater than 0.05 was counted for each strain signal. If any strain showed less than 2 oscillations the particle was rejected. ABC SMC was conducted with population sizes of 10, repeated 255 times yielding a combined final population of 2550 particles.

### 4.7 Random forest classifier model

Using the sci-kit learn (sklearn) python package [38], a random forest classifier was trained using 2000 estimators. The data used consisted of 3750 oscillatory input vectors, and 3750 chaotic input vectors. Training and test datasets were generated with a ratio of 0.5 by random sampling. Figure 6 shows the performance of the classifier model on the test data.

**Figure 6:**
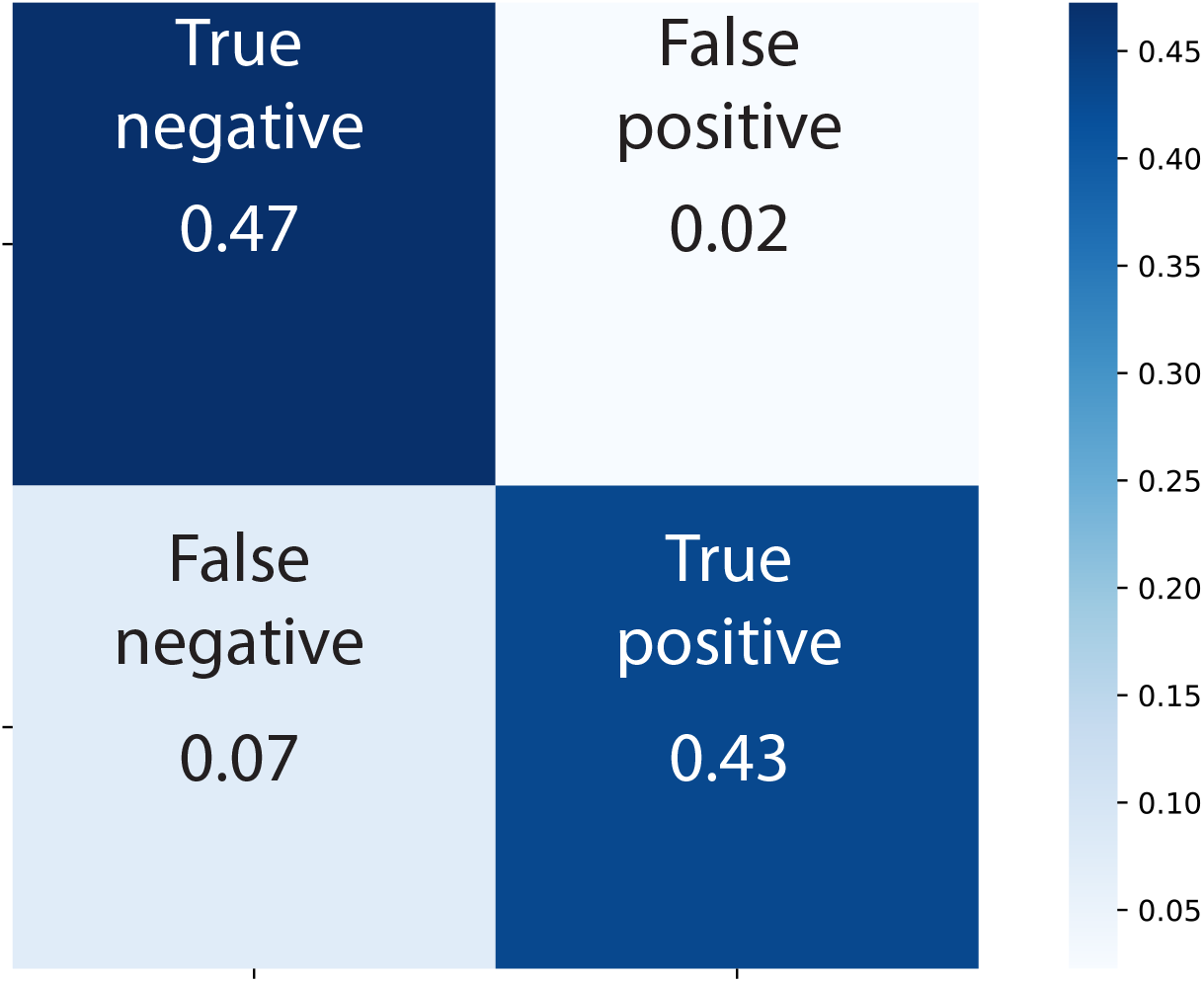
Confusion matrix showing accuracy of random forest classifier on test data

**Algorithm 1:**
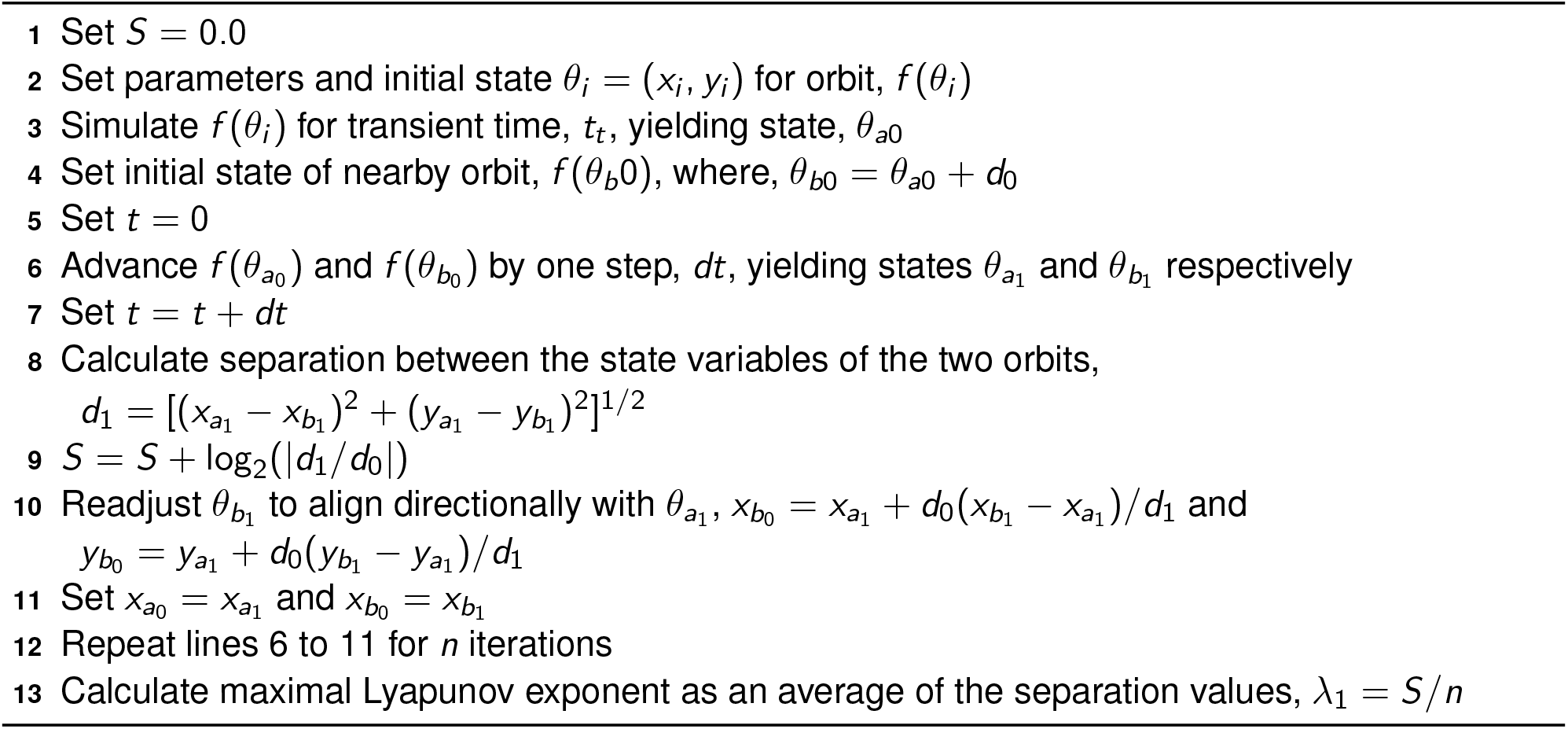
Description of dual-orbit method, demonstrated with two-dimensional system.

### 4.8 Analysis of *m*_850_

*m*_850_ is described by the following equations

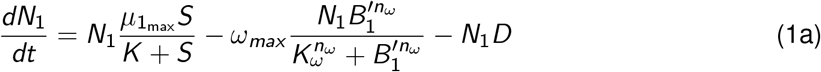

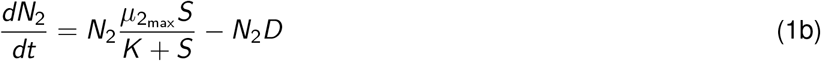

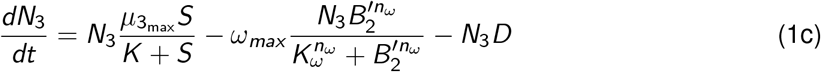

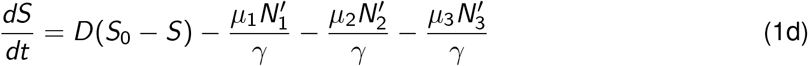

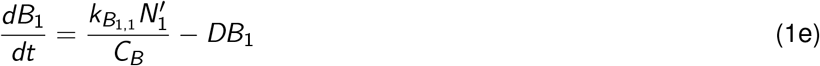

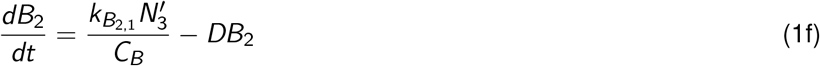

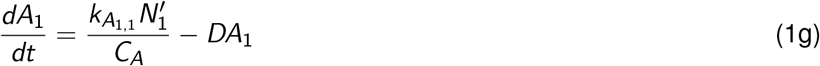

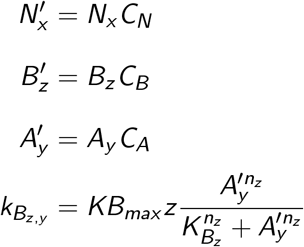

By setting the right hand side of (1) to 0 we find a number of steady states **P** = (*N*_1_, *N*_2_, *N*_3_, *S, B*_1_, *B*_2_, *A*_1_).

#### The trivial steady state

**P**_0_ = (0, 0, 0, *S*_0_, 0, 0, 0). The Jacobian of the linearisation has eigenvalues

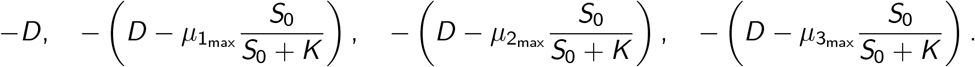

Consequently the trivial steady state always exists and is linearly stable for

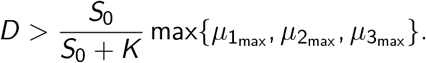

This shows that if the dilution rate is high enough, no strain can survive.

#### One strain only steady states

There are three steady states where only one strain survives, **P**_1_, **P**_2_, **P**_3_. While **P**_2_, and **P**_3_ can be calculated explicitly, **P**_1_ is given implicitly (see below).

We start with **P**_2_:

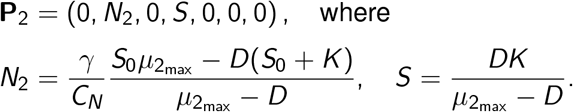

We see that **P**_2_ exists provided

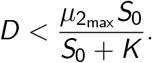

The linearisation at **P**_2_ has eigenvalues

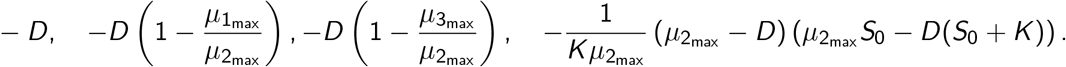

This shows that **P**_2_ exists and is linearly stable if

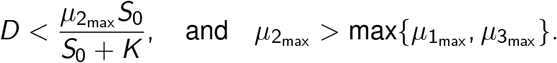

The situation for **P**_3_ is very similar:

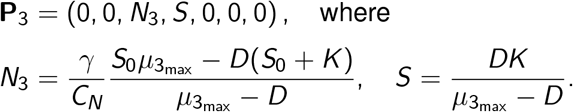

We see that **P**_3_ exists provided

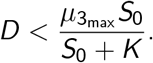

The linearisation at **P**_3_ has eigenvalues

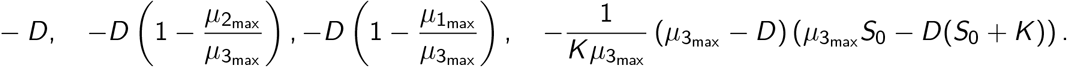

This shows that **P**_3_ exists and is linearly stable if

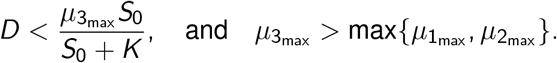

The steady state **P**_1_ = (*N*_1_, 0, 0, *S, B*_1_, 0, *A*_1_) is more complicated and can not be expressed explicitly. Instead it is given as follows: Assume there exists a solution *S* to the following equation

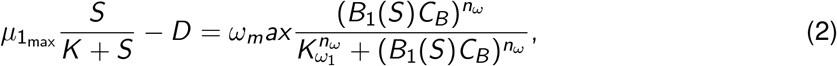

where

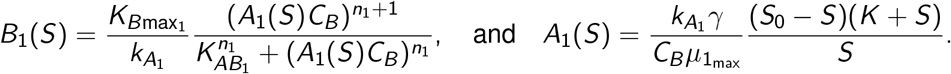

If such a solution *S* exists then *B*_1_ = *B*_1_(*S*), *A*_1_ = *A*_1_(*S*) and 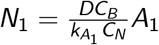.

##### Lemma 1

*There exists a unique steady state* **P**_1_ *if and only if*

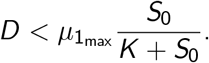

**Proof:** We need the solution to (2) to fulfil *S < S*_0_ in order for *A*_1_ to be positive. We interpret the right and left-hand-sides of (2) as a functions of *S*, denoting them by *R*(*S*) and *L*(*S*) respectively. It is easy to see that *A*_1_(*S*) is a decreasing function of *S, B*_1_(*S*) increases as a function of *A*_1_ and *R*(*S*) is an increasing function of *B*_1_. Consequently the *R*(*S*) is a decreasing function of *S*. We also see that *R*(0) = *ω*_*m*_*ax >* 0 and *R*(*S*_0_) = 0. Further *L*(0) = −*D* and 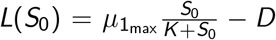 and *L*(*S*) increases as function of *S*. This proves the statement. □.

To summarise, the single-strain survival steady state requires the corresponding maximal growth rate to be large compared to other parameters.

#### The three-strain co-existence steady state

**P**_123_ = (*N*_1_, *N*_2_, *N*_3_, *S, B*_1_, *B*_2_, *A*_1_). From the equation for *N*_2_ we obtain that

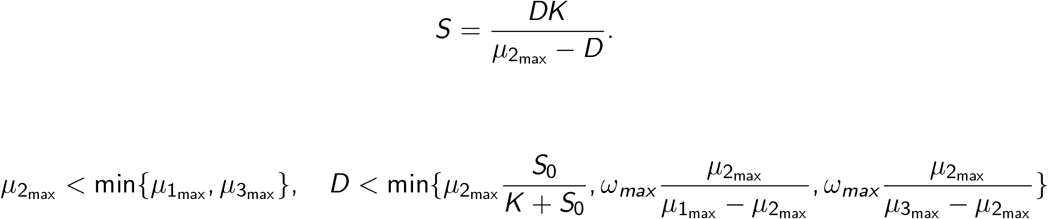

##### Stability

Solved numerically using MATLAB. For each of the 3750 chaotic input vectors we used numerical root finding to calculate **P**_123_, and determined its stability by numerically determining the eigenvalues of the Jacobian. We found **P**_123_ existed for all 3750 input vectors and was stable for 7.8% of them.

**Algorithm 2:**
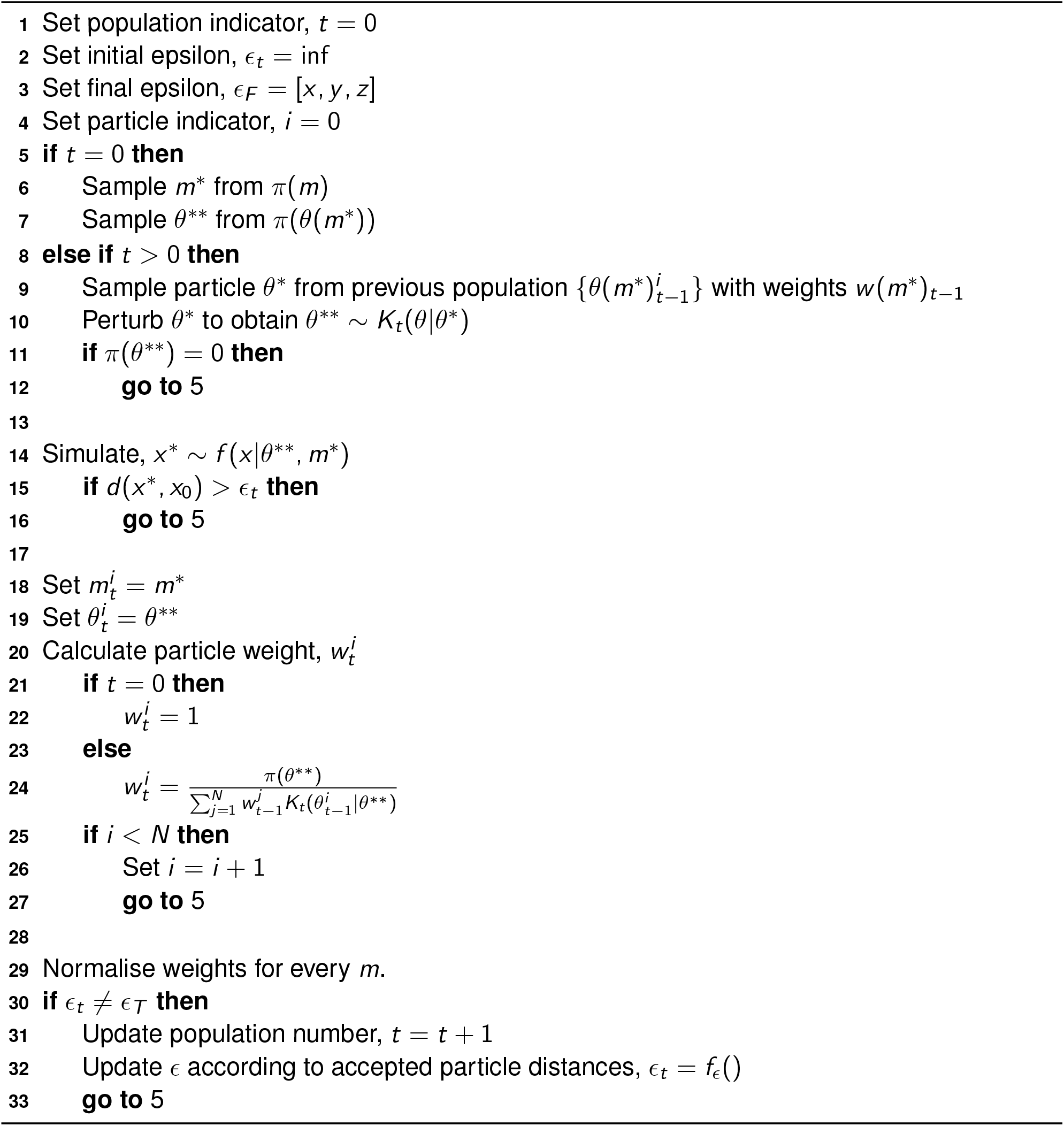
Algorithm for model selection with ABC SMC.

